# Bocaparvovirus, Erythroparvovirus and Tetraparvovirus in New World Monkeys from Central America

**DOI:** 10.1101/588079

**Authors:** A. Chaves-Friedlander, C.N. Ibarra-Cerdena, A.M. López-Pérez, O. Monge, R. Avendaño, H. Ureña-Saborio, M. Chavarría, K. Zaldaña, L. Sánchez, E. Ortíz-Malavassi, G. Suzan, J. Foley, G.A. Gutiérrez-Espeleta

## Abstract

Parvoviruses in the genera Bocaparvovirus (HBoV), Erythroparvovirus (B19) and Tetraparvovirus (PARV4) are the only autonomous parvoviruses known to be associated with human and non-human primates based on studies and clinical cases in humans worldwide and non-human primates in Asia and Africa. Here, the presence of these pathogenic agents was assessed by PCR in blood and feces from 55 howler monkeys, 112 white-face monkeys, 3 squirrel monkeys, and 127 spider monkeys in Costa Rica and El Salvador. Overall, 3.7% of the monkeys had HboV DNA, 0.67% had B19 DNA, and 14.1% had PARV4 DNA, representing the first detection of these viruses in New World monkeys. Sex was significantly associated with the presence of HBoV, males having risk up to nine times compared with females. Captivity was associated with increased prevalence for PARV4 and when all viruses were analyzed together. This work underscores the importance of future research aimed at understanding how these viruses behave in natural environments of the Neotropics, and what variables may favor their presence and transmission.

## INTRODUCTION

The family *Parvoviridae* is a group of small (approx. 5,000 bp) non-enveloped viruses with a linear single-stranded DNA genome (Cotmore et al., 2014). The subfamily *Parvovirinae* infects vertebrates and has eight genera, which are distinguished by their genetic diversity, host range, and pathogenicity (Cotmore et al., 2014, Kailasan et al., 2015). The genera Bocaparvovirus, Erythroparvovirus, and Tetraparvovirus within *Parvovirinae* are the only autonomous parvoviruses associated with human and nonhuman primates (Kailasan et al., 2015). Tetraparvovirus includes human parvovirus 4 (PARV4) and chimpanzee parvovirus 4 (Ch-PARV4) (Kailasan et al., 2015; Sharp et al. 2010), Bocaparvovirus includes human Bocaparvovirus (HBoV 1, 2, 3, and 4), chimpanzee and gorilla bocavirus (GBoV), and Erythroparvovirus encompasses human parvovirus (B19), simian parvovirus (SPV), and macaque parvoviruses (RhMPV and PtMPV) (O’Sullivan et al., 1994; Brown et al., 1995; Kapoor et al., 2010). Parvoviruses may be transmitted through aerosol (e.g. HBoV 1), fecal-oral (HBoV 2, 3, and 4. B19V, PARV4), and respiratory, parenteral and vertical transmission routes (PARV4) (Simmonds et al., 2007; Matthews et al., 2014, Ison et al., 2017; Qiu et al., 2017, Drexler et al. 2012).

Most of what is known about primate-infecting parvoviruses is based on human research, with studies on non-human primates having focused exclusively on Asia and Africa (Kapoor et al., 2010; Sharp et al., 2010, Simon, 2008; Adlhoch et al., 2012). This leaves a gap of information on the genetic diversity and ecology of parvoviruses in New World primates (NWP), species with distinct life histories and ecological dynamics from those already studied. Yet host ecology and virus transmission are important determinants of virus spread and disease impact. Different variables likely influence the prevalence of parvoviruses in primates from different geographic regions (Panning et al. 2010, Fryer et al., 2007; Lucharchaiwong et al., 2008). Host factors associated with foraging habits, metabolic demand, volume of food intake, and management conditions may affect direct and indirect exposure to infectious agents (Matthews et al., 2014; Adlhoch et al., 2012; Fryer et al., 2007; Kesebir et al., 2006; Kumar et al. 2011). The NWP (infraorder Platyrrhini) comprise five families and 19 genera distributed in Central and South America (Rosenberger et al., 2001; Rylands et al., 2012) and among them are species with substantial variation in their life histories and behaviors (e.g. omnivorous vs. herbivorous) (Rosenberger et al., 2001). Although the Old World primates (OWP), infraorder Catarrhini, are evolutionarily distant from the NWP (Perelman et al., 2011), there are social and ecological similarities especially among gregarious species. Among gorillas (*Gorilla* spp.) and chimpanzees (*Pan* spp.), environmental transmission routes for parvovirus transmission seem to predominate (Sharp et al., 2010), which suggests that ground foraging may be a risk factor for omnivorous species. The risk increases in males, due to more interaction with other individuals from their troop and other troops, and higher levels of dispersal than females (Kitchen et al., 2004; Rose, 1994). Attempts at species management can also influence pathogen presence by influencing levels of stress, immunosuppression, interaction with other host species, changes in social habits, and clustering of individuals (Sharp et al., 2017). Although there are no direct comparative studies between free-range and captive OWP, high prevalences to PARV4 and HBoV have been reported in OWP maintained in captivity (Kapoor et al., 2010; Sharp et al., 2010).

The present study aimed to relate biological variables including species, diet, sex, and variables related to management condition including captivity, released from captivity, or free-ranging, to the presence of Bocaparvovirus, Erythroparvovirus and Tetraparvovirus in NWP, using samples collected between 1993 – 2015 in Costa Rica and El Salvador.

## MATERIALS AND METHODS

### Ethics statement

This study was conducted under protocols established by the Institutional Committee for the Care and Use of Animals (Comité Institucional para el Cuidado y Uso de los Animales) of the Universidad de Costa Rica, adhered to the legal requirements of Costa Rica and El Salvador, and adhered to the American Society of Primatologists (ASP) Principles for the Ethical Treatment of Non-Human Primates. Collection permit numbers were MINAET-SINAC-Costa Rica: 042-2012-SINAC, Salvador: MARN-DEV_GVS_AIMA-098-2017.

### Study area and sampling design

Research was carried out using a sample bank collected for a long-term non-human primate genetic variability project. Samples were taken from live monkeys that were captured in the wild, rescue centers, and zoos. Blood samples were collected from free-ranging and captive howler, white-face, spider, and squirrel monkeys from 1993 to 2015 throughout Costa Rica. Fecal samples were collected from spider monkeys in the wild and released from captivity from El Salvador between 2013 and 2015 (Figure 1).

**Figure 1.**
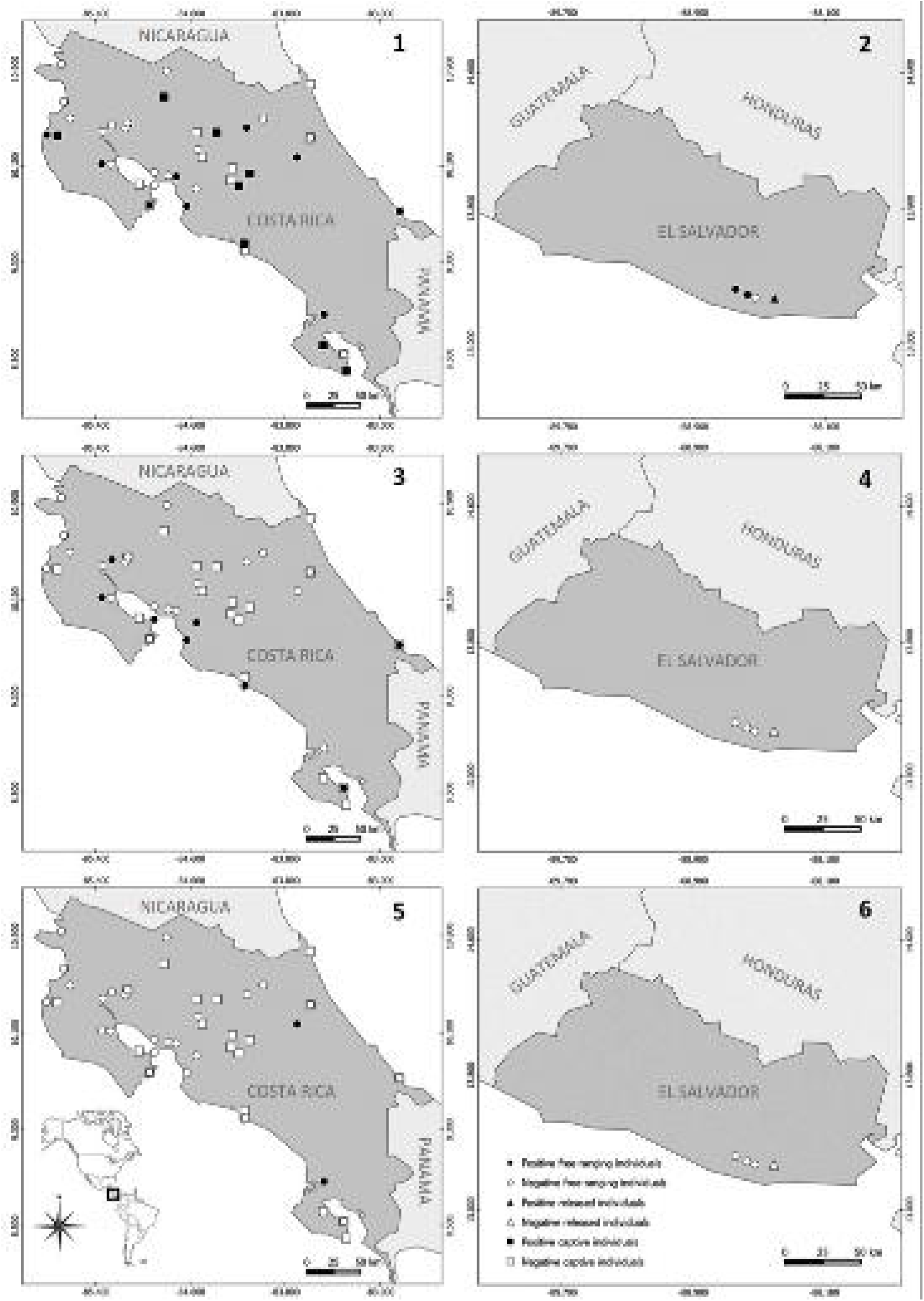
Location of field sites: 1. *Tetraparvovirus* (PARV4) in Costa Rica, 2. *Tetraparvovirus* (PARV4) in El Salvador, 3. *Bocaparvovirus* (HBoV) in Costa Rica, 4. *Bocaparvovirus* (HBoV) in El Salvador, 5. *Erythroparvovirus* (B19) in Costa Rica, and 6. *Erythroparvovirus* (B19) in El Salvador. White circles: free-ranging monkey samples, black circles: free-ranging positives, white triangles: released monkey samples, black triangles: released monkey positives, white squares: captive monkey samples, and black squares: captive monkey positives.

The Costa Rican animals were captured by chemical immobilization with darts (Type P, 1ml, Pneu Dart Inc.), and compressed gas rifle (X-Caliber Gauged CO2, Pneu Dart Inc.) for individuals over long distances, or blowgun for individuals located closely. Anesthetics used were Zoletil 50® (3.3-11 mg/kg), or ketamine (5-20 mg/kg), in combination with xylazine (0.5-2 mg/kg) (Glander et al., 1991; Varela, 2006; West et al., 2007). As soon as the animal was anesthetized, the geographic coordinate was recorded and a 2-4 ml blood sample taken from the femoral, saphenous, or cephalic vein. The samples were collected in tubes with EDTA, and maintained at 4°C until arrival at the laboratory. The monkeys were monitored until awakening from anesthesia and safely released at the capture site. Fresh feces from spider monkeys released from captivity in El Salvador were collected noninvasively into individual sterile vials and maintained at 4°C until arrival to the laboratory. Samples were kept at − 80 °C until analysis.

### Molecular assays

In order to avoid multiple fecal samples from the same individual from El Salvador, the monkey’s identity for each sample was verified by molecular procedures. First, DNA was extracted with the QIAamp DNA Stool Mini kit (Qiagen, Hilden, Germany). Nine pairs of microsatellite loci [AP74, Ceb120, Ceb 121, D8S260, Leon2, LL1110, SB38, P2BH6 and D8S165] were analyzed according to Sánchez and Zaldaña^38^. With the GIMLET 1.3.3 program^39^, the probability of identical genotypes was calculated (probability> 90%). Samples with missing data for more than two loci and samples with repeated genotypes were discarded.

Total DNA was extracted from blood samples using the DNeasy® Blood & Tissue Kit (Qiagen, Hilden, Germany) and from all feces samples using the QIAamp DNA Stool Mini kit (Qiagen, Hilden, Germany), according to the manufacturer’s instructions. The DNA samples were quantified using a NanoDrop 2000 spectrophotometer (ThermoFisher, USA). Polymerase chain reaction (PCR) was carried out as described previously^8^. PCR products were visualized on 1% agarose gels stained with Gel Red (Biotium Inc., Hayward, California, USA). DNA from *Bocaparvovirus, Erytroparvovirus*, and PARV4 donated by the Roslin Institute, University of Edinburgh was used for positive controls. Nuclease-free water was used as negative control.

PCR products were sequenced using BigDye Terminator v3.1 cycle sequencing kit and cleaned with BigDye XTerminator Purification kit (Applied Biosystems, Foster City, California, USA). Products were analyzed on a 3130xl Genetic Analyzer (Applied Biosystems). Sequence alignments were performed using Clustal W (Larkin et al., 2007). Phylogenetic and molecular evolutionary analyses were conducted using the distance matrix method (maximum likelihood algorithm) and the nucleotide distance measure based on the best model per data set; the complete deletion option within MEGA X (Kumar et al., 2018) was also used. Data sets were bootstrapped based on 1000 resamplings of the original data set to produce a majority rule consensus tree. Sequences were deposited in GenBank (accessions MF476926 to MF476982). Host species for MF476960 and MF476962 could not be identified and are not included in further analyses.

### Statistical analysis

Logistic regression was used to evaluate the possible associations between PCR-positivity (one model considering infection regardless of virus species, and separate models for each virus species) and primate species, diet (omnivorous or herbivorous), and management condition (captive, released from captivity, or free-ranging). Stepwise model selection using AIC was performed using the function stepAIC of the package MASS ver 7.3-50 (https://cran.r-proiect.org/web/packages/MASS/MASS.pdf) in R ver 3.4.2 (R. Core Team, 2013). Because 33% of the host individuals were not sexed, we did not include this variable in the models, but its effect was tested separately using odd ratios with the 95% interval of confidence. Analyses with P < 0.05 were considered significant.

## RESULTS

A total of 297 individual non-human primates were sampled (243 from Costa Rica, and 54 from El Salvador), of which 55 were howler monkeys (*Alouatta palliata*), 112 were white-face monkeys (*Cebus imitator*), 3 were squirrel monkeys (*Saimiri oerstedii*), and 127 were spider monkeys (*Ateles geoffroyi*) (73 in Costa Rica, and 54 in El Salvador). Of these, 198 were free-ranging, 89 were kept in captivity, and 10 animals were being released from captivity. In Costa Rica, only free-ranging and individuals kept in captivity were analyzed, and in El Salvador, free-ranging individuals and those being released from captivity were analyzed (Fig. 1, Table 1).

**Table 1.** Number of monkeys of each species, sex, diet, and management status from which Tetraparvovirus (PARV4) Bocaparvovirus (HBoV), and Erythroparvovirus (B19) PCR-positive samples were collected in Costa Rica and El Salvador.

|  | Number of PCR-Positive Monkeys/Number Tested |  |  |
| --- | --- | --- | --- |
|  | Species |  |  |
|  | PARV4 | HBoV | B19 |
| *Alouatta palliata* | 6/55 | 4/55 | 1/55 |
| *Cebus imitator* | 13/112 | 7/112 | 1/112 |
| *Ateles geoffroyi* | 23/127 | 0/127 | 0/127 |
| *Saimiri oerstedii* | 0/3 | 0/3 | 0/3 |
| **Total** | **42/297** | **11/297** | **2/297** |

|  | Sex*<sup>1</sup> |  |  |  |  |  |
| --- | --- | --- | --- | --- | --- | --- |
|  | Male |  |  | Female |  |  |
|  | PARV4 | HBoV | B19 | PARV4 | HBoV | B19 |
| *Alouatta palliata* | 3/28 | 4/28 | 0/28 | 2/23 | 0/23 | 1/23 |
| *Cebus imitator* | 5/54 | 5/54 | 1/54 | 8/36 | 1/36 | 0/36 |
| *Ateles geoffroyi* | 10/29 | 0/29 | 0/29 | 9/44 | 0/44 | 0/44 |
| *Saimiri oerstedii* | ND | ND | ND | ND | ND | ND |
| **Total** | **18/111** | **9/111** | **1/111** | **19/103** | **1/103** | **1/103** |

|  | Diet |  |  |  |  |  |
| --- | --- | --- | --- | --- | --- | --- |
|  | Herbivorous |  |  | Carnivorous |  |  |
|  | PARV4 | HBoV | B19 | PARV4 | HBoV | B19 |
| *Alouatta palliata* & *Ateles geoffroyi* | 29/182 | 4/182 | 1/182 | N/A | N/A | N/A |
| *Cebus imitator* & *Saimiri oerstedii* | N/A | N/A | N/A | 13/115 | 7/115 | 1/115 |
| **Total** | **29/182** | **4/182** | **1/182** | **13/115** | **7/115** | **1/115** |

|  | Management |  |  |  |  |  |  |  |  |
| --- | --- | --- | --- | --- | --- | --- | --- | --- | --- |
|  | Free-ranging |  |  | Released |  |  | Captivity |  |  |
|  | PARV4 | HBoV | B19 | PARV4 | HBoV | B19 | PARV4 | HBoV | B19 |
| *Alouatta palliata* | 5/52 | 4/52*<sup>2</sup> | 1/52*<sup>3</sup> | N/A | N/A | N/A | 1/3 | 0/3 | 0/3 |
| *Cebus imitator* | 10/96 | 7/96 | 1/96*<sup>4</sup> | N/A | N/A | N/A | 3/16 | 0/16 | 0/16 |
| *Ateles geoffroyi* | 3/47 | 0/47 | 0/47 | 1/10 | 0/10 | 0/10 | 19/70 | 0/70 | 0/70 |
| *Saimiri oerstedii* | 0/3 | 0/3 | 0/3 | N/A | N/A | N/A |  |  |  |
| **Total** | **18/198** | **11/198** | **2/198** | **1/10** | **0/10** | **0/10** | **23/89** | **0/89** | **0/89** |

\*<sup>1</sup> The missing data were individuals with no determined sex data
\*<sup>2</sup> Co-infection of HBoV and PARV4 was detected in one individual
\*<sup>3</sup> Co-infection of B19 and PARV4 was detected in one individual
\*<sup>4</sup> Co-infection of B19 and PARV4 was detected in one individual

### Primate parvoviruses prevalence

All PCR-positive spider monkeys (23/127, 18.1%) were infected with PARV4, and the 3 squirrel monkey samples were PCR-negative. Nine of 55 (16.4%) howler monkeys were PCR-positive, of which 6 (10.9%) represented PARV4, 4 (7.3%) were HBoV, and 1 (1.8%) was B19. A total of 17.9% (20/112) white-face monkeys were positive to parvovirus, 11.6% (13/112) to PARV4, 6.2% (7/112) to HBoV, and 0.9% (1/112) to B19. One howler monkey was simultaneously infected with PARV4 and HBoV, and one howler monkey and one white-face monkey were simultaneously infected with PARV4 and B19. The total prevalence for parvovirus infection among all monkeys was 17.5% (52/297).

### Genetic diversity of primate parvoviruses

Overall, of the 52 parvovirus PCR-positive monkeys, 49 (94.2%) originated from Costa Rica and 3 (5.8%) came from El Salvador. Forty-two 42 (14.1%) of the samples represented PARV4 (39 from Costa Rica and 3 from El Salvador), 11 (3.7%) were HBoV, and 2 (0.67%) were B19 (Table 1).

Phylogenetic analysis of the 42 PARV4 sequences showed a total of 10 genetic variants with a similarity of 99.66 to 98.64% among them (Figure 2). All Central American PARV4 strains were in the “genotype 1” clade, with a similarity of 98.99% to the DQ873386 sequence previously reported from a human from the USA (Fryer et al. 2007). There were six HBoV variants, with 99.71 to 99.44% among them, all within genotype 3, and with 98.31% similarity to sequence FJ973562 from a human from Tunisia (Kapoor et al 2010) (Figure 3). The two Costa Rican B19 variants shared 98.10% similarity and grouped within “genotype 1”, with a similarity of 98.34% to sequence EU478553 from a human from the United Kingdom (Norja et al 2008) (Figure 4).

**Figure 2.**
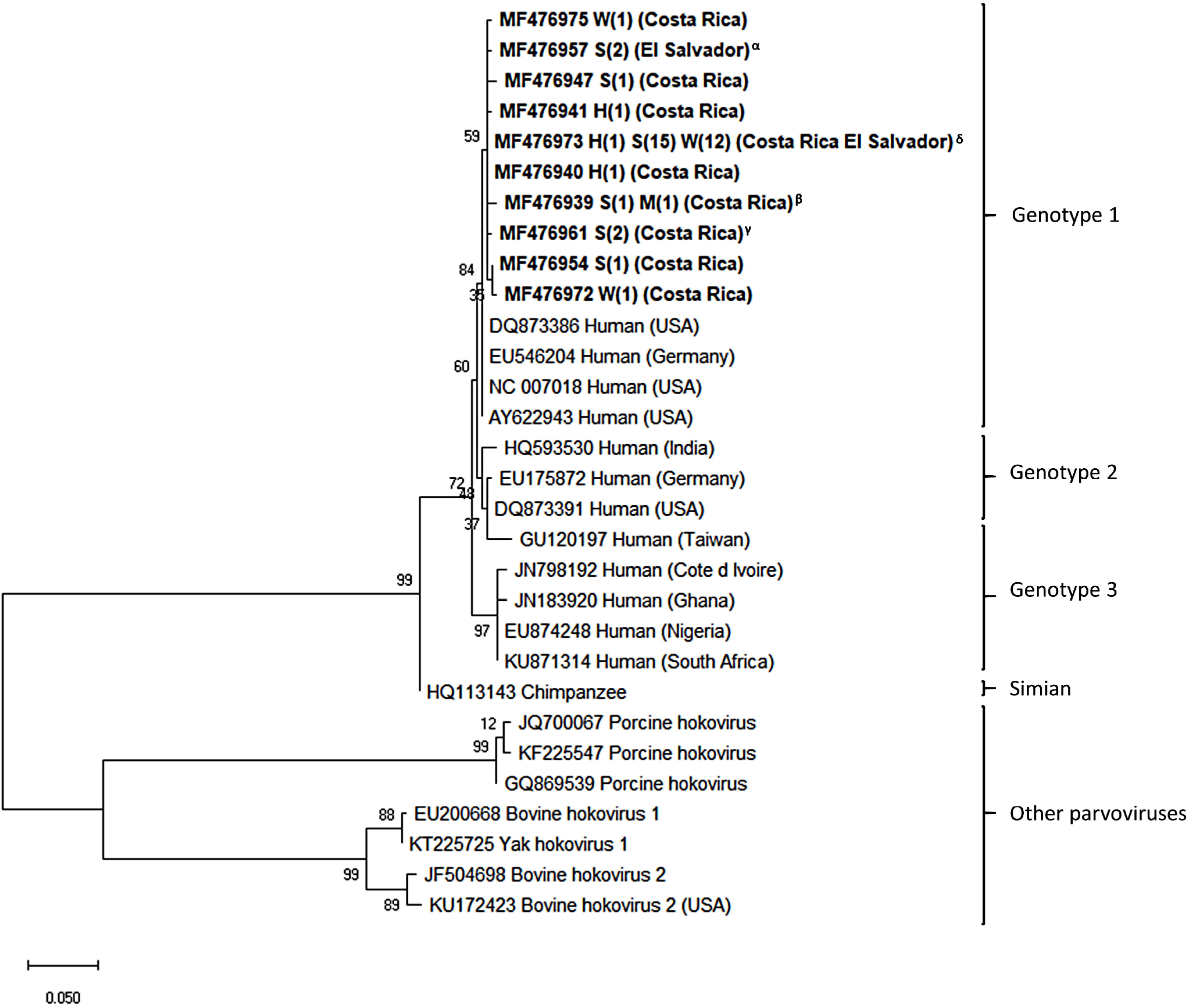
Maximum-likelihood (ML) tree of tetraparvovirus 4 based on VP1 partial sequences. ML bootstrap proportion greater than 50% is presented at the nodes. Capital letters refer to host species (H, howler monkey; S, spider monkey; W, white-headed capuchin). Numbers in parentheses are the number of sequences obtained from monkeys. Identical sequences are clustered to representative accessions: ^α^:MF476957; ^β^: MF476939; ^γ^: MF476948; ^δ^: MF476973 (Full list of sequences: suppl. table 1). The tree was rooted to other tetraparvoviruses: ungulate tetraparvovirus 1 (bovine hokovirus 1 & 2) and ungulate tetraparvovirus 2 (porcine hokovirus).

**Figure 3.**
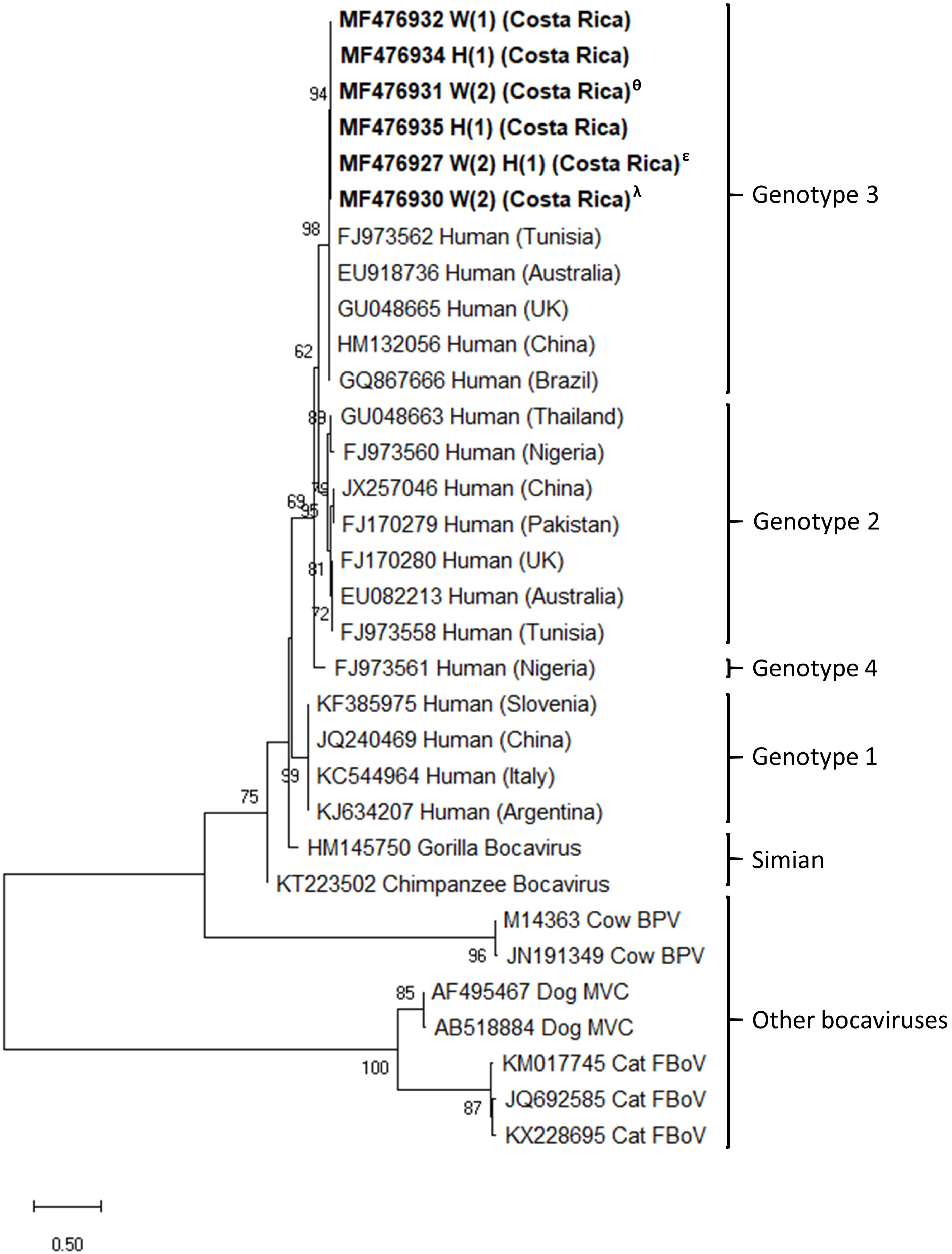
Maximum-likelihood (ML) tree of bocavirus based on VP1 & VP2 partial genes sequences. ML bootstrap proportion greater than 50% is presented at the nodes. Genotypes from this study are in bold. Capital letters refer to host species (H, howler monkey; W, white-face monkey). Numbers in parentheses are the number of sequences obtained from monkeys. Identical sequences are clustered to representative accessions:^ε^: MF476927; ^θ^: MF476931; ^λ^: MF476930 (full list of sequences: suppl. table 1). The tree was rooted to other bocaviruses: carnivore bocavirus 1 (dog), carnivore bocavirus 3 (feline) & Ungulate bocavirus 1 (bovine).

**Figure 4.**
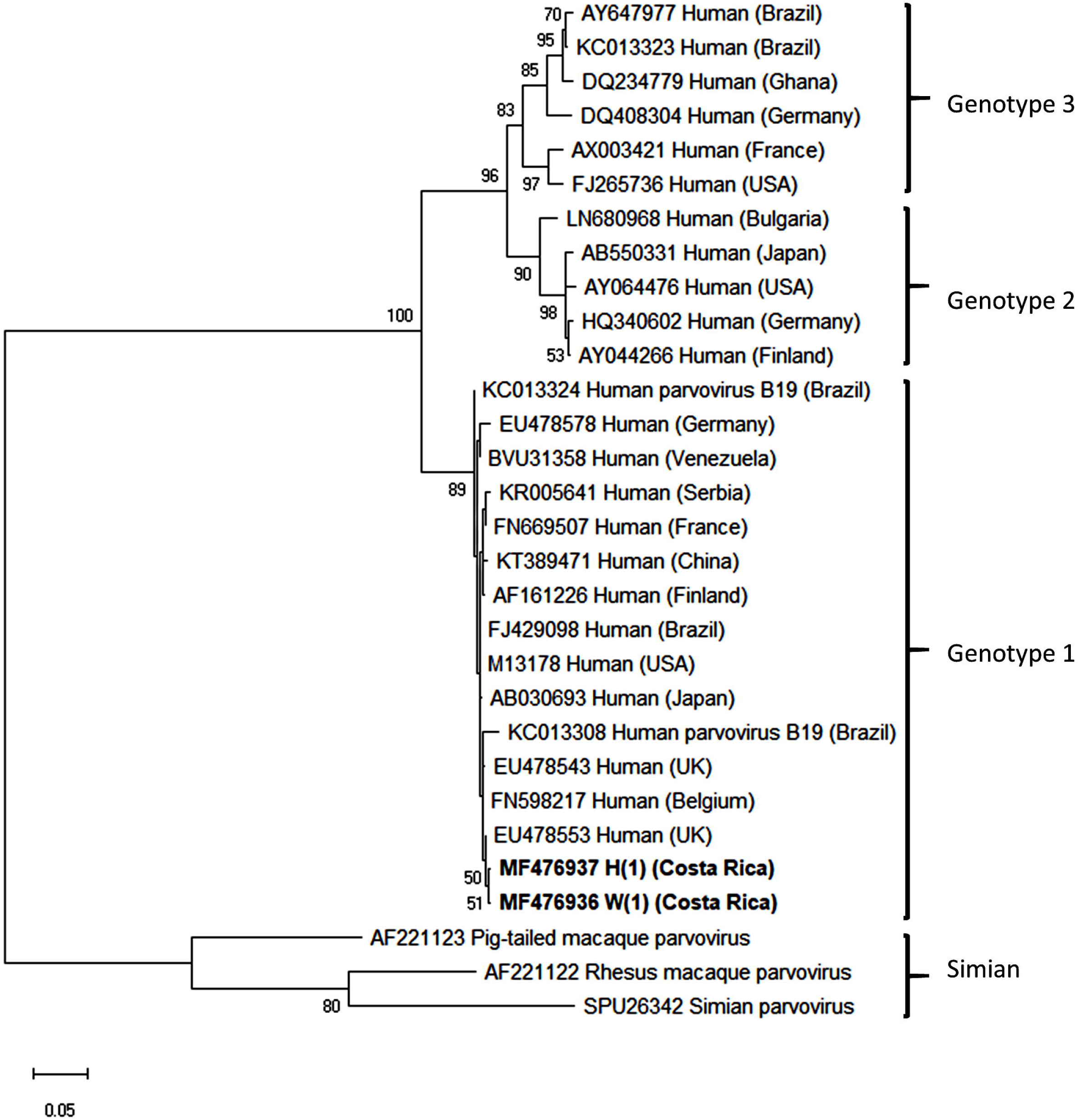
Maximum-likelihood (ML) tree of erythrovirus based on VP1 & VP2 partial genes sequences. ML bootstrap proportion greater than 50% is presented at the nodes. Genotypes from this study are in bold. Capital letters refer to host species (H, howler monkey; W, white-face monkey). Numbers in parentheses are the number of sequences obtained from monkeys. The tree was rooted to simian erythroviruses.

### Biological and management variables

Among candidate risk factors for PCR-positive test results, management and sex were significant predictors. Management was only marginally important in the model “general infection” (all parvoviruses) (Deviance=5.7; DF=2; p=0.06), but highly important in the PARV4 model (Deviance=12.88; DF=2; p=0.002), where being kept in captivity resulted in a 2.42 fold increase in risk compared with free ranging monkeys. There was no effect of sex on PCR-positivity when all viruses were analyzed together (male to female odds ratio: 1.27; 95% CI: 0.66 to 2.45), PARV4 (male to female odds ratio: 0.86; 95% CI: 0.42-1.74) or B19 (male to female odds ratio: 0.93; 95% CI: 0.06-15.02). However, males were much more likely to be infected with HBoV than females (odds ratio: 9; 95% CI: 1.12 to 72.34).

## DISCUSSION

Little is known of the parvoviruses present in non-human primates, their transmission routes, and factors associated with their presence or prevalence, with most previous investigations being phylogenetic or serological, comparing parvoviruses in human and non-human primates especially in Asia and Africa (Kapoor et al., 2010; Sharp et al., 2010; Simon, 2008; Adlhoch et al., 2012). This study is the first molecular characterization of circulating genotypes of PARV4, HBoV and B19 in NWP, documenting genotype 1 of PARV4, genotype 3 of HBoV, and genotype 1 of B19 in NWP in Central America. PARV4 is classified into three genotypes, of which 1 and 2 are predominant in Europe and the United States (Manning et al., 2007; Nguyen et al., 1999) and 3 is mostly found in Africa (Parsyan et al., 2007). The close genetic relationship between the NWP and North American human PARV4 sequences suggests possible zoonoses, in contrast to PARV4 virus in North America and Europe which are specifically human (Matthews et al. 2014), and African and Asian PARV4 viruses which feature separate non-human and human clades even where there is frequent inter-species contact (Sharp et al., 2010; Adlhoch et al., 2012).

Similarly with PARV, both HBoV and B19 clustered together with sequences reported in humans (Guido et al., 2016). It has been suggested that chimpanzee bocavirus and HBoV 1 diverged from a common ancestor, HBoV 4 separated independently from the great ape’s bocavirus 200 to 300 years ago, and recombination of HBoV 1 and HBoV 4 was the source of HBoV 2 and HBoV 3 (Kapoor et al., 2010; Babkin et al., 2013). There are three different genotypes of B19, of which genotype 1 (subtypes 1a and 1b) is the most prevalent worldwide, genotype 2 is only detected in patients born before 1973 in European countries and North and South America, and genotype 3 (subtypes 3a and 3b) predominates in West Africa (Servant et al., 2002). Clustering of human and non-human primate HBoV and B19 have been described before (Servant et al., 2002; Manning et al., 2007; Kumakamba et al., 2018), suggesting cross species transmission or that the dissemination of the parvovirus genotypes has been a relatively recent phenomenon.

Among the NWP in the present study, the highest parvovirus prevalence occurred for PARV4 in free-ranging individuals, and PARV4 was also the only parvovirus detected in captive and released monkeys. High PARV4 seroprevalences have also been reported for gorillas (*Gorilla gorilla)* (18%), chimpanzees (*Pan troglodytes*) (2-60%) (Sharp et al., 2010), red colobus monkeys (*Piliocolobus badius*) (23%), black-and-white colobus monkeys (*Colobus polykomos*) (13%), and baboons (*Papio anubis*) (5%) (Sharp et al., 2010; Adlhoch et al., 2012). Although an environmental source of infection is suspected for non-human primates (Adlhoch et al., 2012), transmission routes of PARV4 in non-human primates are not known. Several and different transmission routes are possible even for African humans, which might be even more than those reported for humans from North America and Europe (Matthews et al., 2014; Panning et al., 2010; Fryer et al., 2007; Lucharchaiwong et al., 2008). Despite in West Africa, where opportunities for cross-species transmission abound, Parv4-like viruses are reported to be species-specific (Adlhoch et al., 2012), suggesting that the risk of transmission from NWP to humans is low. However, it is suggested that the genotypes of PARV4 may represent different zoonotic transmissions (Manning et al., 2007; Sharp et al., 2010). Captivity was a significant risk factor for PARV4 in howler, white-face capuchin and spider monkeys. PARV4 risk is elevated in immunosuppressed humans and other animals (Matthews et al., 2014; Fryer et al., 2007). Animals in captivity may experience increased stress (Morgan & Tromborg 2007), and infectious agents are known to increase in prevalence in captive wildlife (Capitanio 2012). Captive environments also facilitate direct and indirect contact among individuals, troops or even different monkey species including through common waterways, food handling, proximity of traps, and transfer of animals into captive centers, which may favor PARV4 transmission (Drexler et al., 2012; Panning et al., 2010).

Free-ranging monkeys were also infected with HBoV (Table 1). In Africa, seroprevalence as high as 73% in chimpanzees and (36.3%) (Jones et al., 2005) in gorillas have been identified (Sharp et al., 2010). With high PCR-prevalences of HBoV 1, 2, 3, and 4 worldwide (Aderson et al., 2007; Manteufel, & Truyen 2008; Allander, 2008), it has recently been suggested that HBoV is much more widely dispersed than previously known (Kapoor et al., 2010). Human bocaviruses with very high similarity to HBoV 2 and 3 were recently described from OWP in the Democratic Republic of the Congo, suggesting possible transmission between humans and non-human primates (Kumakamba et al., 2018). In the present study, sex of the host species was significantly associated with HBoV. This virus is typically transmitted by fecal-oral and aerosol routes (Kesebir et al., 2006; Kumar et al., 2011), possibly after environmental contamination. Male monkeys typically have higher levels of social interaction and dispersal than females; for example, males usually change troop when they reach adulthood while females stay in the same troop (Kitchen, 2004; Fedigan & Baxter, 1984). Male and female diets may differ as well; for example the white face capuchin monkey male feeds on larger prey than females, covers greater distance, and spends more time foraging (Rose, 1994). All of the factors may increase risk of contact with contaminated environmental sources of HBoV in male NWPs.

B19 (0.6%) was the least abundant parvovirus, as was found previously (Sharp et al. 2010), with 8% seroprevalence in chimpanzees and 27% in gorillas, with all of these OWP all showed negative results OWPs being negative by PCR. Persistent or latent B19 infections are common (Young, & Brown 2004) and may be difficult to detect, such as simian parvovirus (SPV), pig-tailed macaque parvovirus (PtPV), and rhesus parvovirus (RhPV) (O’Sullivan et al., 1994; Green et al., 2000; Simon 2008). We also detected co-infection of PARV4 with HBoV in one and B19 in two free-ranging individuals. It would be of interest to investigate cases of parvovirus co-infection, as this is consistent with animals being immunosuppressed or having anemia and other health problems (Simon, 2008; Fryer et al., 2007).

The detection for the first time of PARV4 in blood and fecal samples in NWP, and of HBoV and B19 in blood samples, is a pioneering line of investigation to understand the way in which these viruses behave in natural environments in the Neotropics, and allows for comparisons with characteristics of these viruses from other continents in both free-range and captive primates (Kapoor et al., 2010; Sharp et al., 2010; Adlhoch et al., 2012). It would be valuable to continue to perform monitoring and conduct longitudinal studies in NWP populations to document the sources and phenology for these viral infections. Other viral species may yet be discovered in NWP as well which could impact conservation of Neotropical primates (Sharp et al., 2010; Simon, 2008). Moreover, with increasing evidence for shared non-human primate and human viral transmission, the risks of zoonoses should be taken into account in situations of interaction between non-human primates and humans, such as captivity centers.

## Acknowledgments

We are grateful to all contributors who assisted with sampling endeavors, especially Mauricio Jimenez, María Isabel DiMari, Sofia Bernal, Genuar Nuñez, Natalia Valverde and Mauricio Losilla. We thank Dr. Colin P. Sharp at the Roslin Institute, University of Edinburgh for the positive parvoviruses controls. We would also like to thank the National System of Conservation Areas (SINAC), the Ministry of Environment and Energy (MINAE) of Costa Rica and the Ministry of Environment and Natural Resources of El Salvador (MARENA).

